# CMVAE: Contrastive Supervision Improves Single-Omic Latent Manifold for Breast Cancer Survival Prognosis

**DOI:** 10.64898/2026.10.30.763085

**Authors:** Akash Singh, Adrian Kohut, John Michael Templeton, Anowarul Kabir

## Abstract

High-dimensional omics data are vital for forecasting patient prognosis in oncology. While deep multi-omics learning methods are increasingly popular, they suffer from data missingness and low patient overlap across modalities. Consequently, advancing single-omic representation learning remains a critical yet challenging research problem. Standard generative dimensionality reduction methods rely on reconstruction objectives that optimize the global manifold topology; however, overlook downstream clinical outcomes. To address these limitations, we introduce a Contrastive Masked Denoising Variational Autoencoder (CMVAE), a self-supervised dual-optimization framework designed to map single-omic information into a clinical outcome-aware latent space. A stochastic feature masking strategy corrupts the input omic profiles, and CMVAE is optimized to simultaneously infer missing molecular data and contrastive alignment of patient survival trajectories. We evaluate CMVAE using leak-proof five-fold stratified cross-validation on the TCGA Breast Invasive Carcinoma (BRCA) cohort. The method outperforms traditional unsupervised methods, achieving an AUROC of 0.7397 ± 0.0239 on patient prognosis. Further analysis demonstrates a highly significant (*p* < 0.0001) risk stratification among low vs. high risk groups. Beyond comparing within single-omic methods, CMVAE outperforms six state-of-the-art multi-omics survival baselines by a minimum of 3%^1^.

**Availability:** All codes including data preparation scripts, training and validation recipes, and experimental configurations are available at: https://github.com/kabir-ai2bio-lab/CMVAE.

## I. Introduction

High-throughput omics technologies [1]–[3] provide a comprehensive characterization of biological mechanisms of a patient at different layers, including genomics, epigenomics, transcriptomics, proteomics and post-transcriptional regulators. Identifying robust molecular signatures from these diverse layers is critical in precision oncology. In recent years, literature has focused on multiomics integration workflows to capture deep phenotypic variations [4]–[7] However, a major bottleneck of multi-omics modeling remains data missingness and baseline scarcity: multi-omic cohorts often suffer from low sample overlap across all modalities [7], [8]. For example, LinkedOmics TCGA-BRCA cohort [9], [10] contains clinical information for 1,097 samples, while only 643 samples have complete data across four omics modalities. Therefore, we ask: *can we learn a robust single-omic representation to accurately forecast prognosis and stratification of critical patients?*

We hypothesize that high-dimensional single-omic profiles may contain highly localized rich biological signals essential for predicting patient clinical outcomes. When tested, standard unsupervised deep generative frameworks, i.e. Autoencoders (AEs) and Variational Autoencoders (VAEs), achieve sub-optimal performance to isolate these boundaries (see Section IV), contrary to the popularity of AEs/VAEs in multiomics integration for patient prognosis or stratification [11]– [15]. We rationalize that standard AE/VAE architectures optimize an unsupervised reconstruction objective [16]–[18] that might learn the global manifold topology, but overlook the localized molecular signatures relevant to downstream clinical outcomes.

To address these limitations, we introduce a **Contrastive Masked Denoising Variational Autoencoder (CMVAE)**, a self-supervised framework designed to bridge representation learning with downstream prognostic utility. The model first applies a stochastic feature masking strategy to corrupt high-dimensional input omic profiles. It forces the model to learn robust, noise-invariant molecular features. CMVAE is then optimized via a joint loss function that simultaneously reconstructs missing molecular data and enforces a supervised contrastive objective. This contrastive component explicitly clusters patients with similar survival criteria while separating divergent clinical trajectories within the latent space. By unifying generative reconstruction with distance-based contrastive regularization, this dual-optimization framework maps single-omic information into a clinical outcome-aware latent space.

We validate our framework across four distinct omics modalities on BRCA patient outcomes: mRNA (transcriptomics), DNA methylation (epigenomics), miRNA (post-transcriptional regulators), and copy number variations (CNV) (genomics). To guaranty an unbiased, leak-proof estimate of model generalizability under severe survival class imbalances, all data preprocessing, imputation, scaling, and oversampling are isolated strictly within a stratified five-fold cross-validation scheme. Our empirical findings demonstrate that CMVAE universally stabilizes and optimizes feature extraction across four distinct modalities. In high-density transcriptomic (mRNA) data, CMVAE outperforms linear and non-linear dimensionality reduction methods in extracting meaningful representation for classifying short-vs. long-term BRCA survival (STS/LTS) patients. It achieves a peak performance of 0.7397 ± 0.0239 AUROC in prognosis. Kaplan-Meier (KM) survivability analysis further shows that CMVAE maps two risk strata into a highly structured latent space with distinct clinical boundaries (*p <* 0.0001). Beyond binary classification, out-of-fold risk scores evaluated via Harrell’s Concordance Index (C-index) [19] reveal a stable survival-ranking signature (0.688 ± 0.031) across all folds. When comparing with multiomics survival approaches, CMVAE outperforms six state-of-the-art methods, including three deep-learning based models, by a minimum of 3%.

## II. Methodology

### A. Leak-proof Data Preprocessing, Training & Validation

To ensure an unbiased evaluation of model generalization and prevent data leakage, all data manipulation steps, such as preprocessing, feature filtering and scaling, representation learning, oversampling, and classifier training, are executed strictly within a fold-specific pipeline. We adopt the following leak-proof steps in all experiments.

For each omic modality, we utilize a five-fold stratified cross-validation with an approximate 80/20 train/validation split. Within each cross-validation loop, the preprocessing pipeline is fitted exclusively on the labeled training partition and then applied to the corresponding validation partition. Since each omic modality has different feature density, missingness pattern, and dimensionality, modality-specific feature filtering thresholds are used. In general, features with excessive missingness are removed, and a fixed number of highly variable features is selected based on variance computed from the training fold. Remaining missing values are handled through the fold-specific imputation strategy. Feature magnitudes are then normalized using *z*-score standardization.

For CMVAE representation learning, confidently labeled training samples are used for both reconstruction and supervised contrastive learning. Ambiguous samples, defined as patients censored before the five-year survival threshold, do not contribute to supervised contrastive learning, classifier training and evaluation. However, since censored omics profiles still contain biologically meaningful structure, they are allowed to contribute only to the reconstruction component of the CMVAE training.

To mitigate class imbalance, oversampling is applied exclusively to the training partition after pre-processing and latent representation extraction. The validation fold remains entirely unmanipulated and preserves its original class distribution. These steps ensure that validation samples do not influence feature filtering, imputation, scaling, representation learning, oversampling, or classifier fitting, thereby providing an unbiased estimate of model generalization performance.

### B. CMVAE: Contrastive Masked Denoising Variational Autoencoder

The proposed CMVAE adopts a standard VAE architecture to learn survival-aware latent representations from high-dimensional single-omic profiles. CMVAE combines two learning signals. First, stochastic feature masking is used to corrupt the input omic profile, forcing the model to infer missing molecular information from the remaining observed features. Second, a supervised contrastive objective is added to organize the latent space according to the clinical outcomes of the patients. Fig. 1 illustrates the overview of the CMVAE pipeline.

**Fig. 1.**
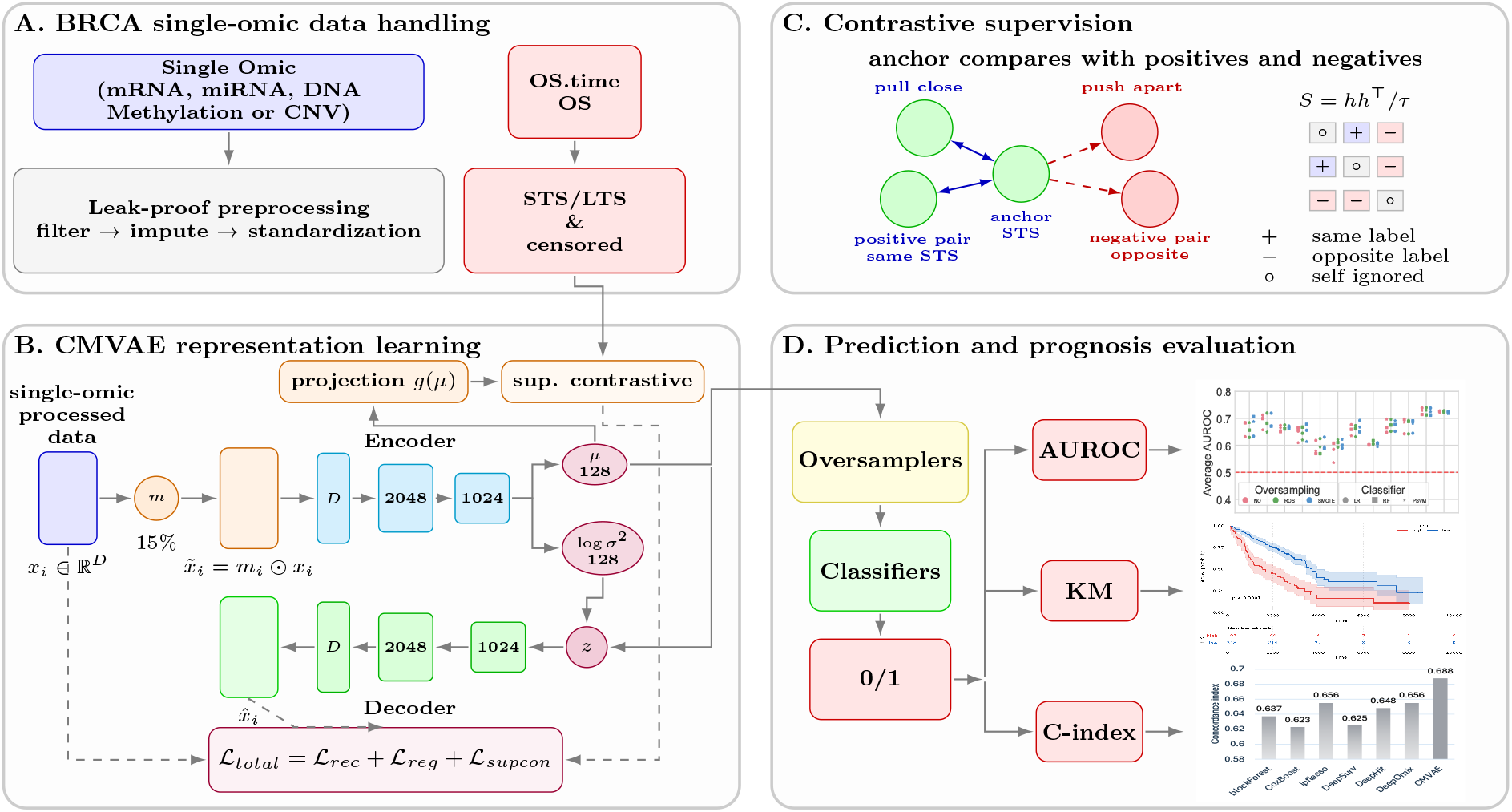
CMVAE pipeline overview. (A) Each omic modality is processed independently through a leak-proof fold-specific preprocessing and aligned with sample labels. (B) The CMVAE representation learning module maps the corrupted single-omic profile into a clinical outcome-aware latent space by simultaneously optimizing to infer missing molecular data and contrastive alignment of patient survival trajectories. (C) A contrastive supervision mechanism. (D) The learned patient embeddings are evaluated onto patient prognosis and survival stratification tasks. Sample analyses (AUROC, Kaplan-Meier (KM) survivability and C-index comparison) are illustrated.

#### 1) Model Architecture: Variational Autoencoder (VAE)

Let *x*_*i*_ ∈ ℝ^*D*^ denotes the preprocessed omics vector for patient *i*, where *D* is the number of features (i.e. genes) for the corresponding omic modality. The encoder consists of two fully connected layers to extract non-linear combination of features. Each layer is followed by a batch normalization (BN) [20] and non-linear ReLU activation. We use BN instead of layer normalization [21] because our inputs are fixed gene-wise omics profiles, and BN normalizes activations across mini-batch samples while retaining a feature-wise representation structure. The encoder then maps the hidden representation into a latent mean *µ*_*ϕ*_(*x*) and a latent log-variance 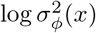. Together, these parameterize the approximate the posterior distribution:

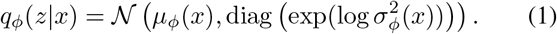

During training, latent sampling is performed through the standard reparameterization trick [22]:

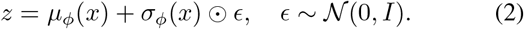

The decoder maps the latent vector back to the original input feature space using fully connected layers, followed by a linear output layer of size *D*. A linear decoder output is used because the preprocessed omic values are continuous-valued. At inference time, the stochastic sampling step is omitted, and the deterministic posterior mean *µ*_*ϕ*_(*x*) is used as the final patient-level latent embedding.

#### 2) Masking Strategy

CMVAE uses stochastic element-wise feature masking before encoding the input omics profiles. For each sample *x*_*i*_, a binary mask *m*_*i*_ ∈ {0, 1}^*D*^ is sampled independently, where each feature is retained with probability 1 − *ρ*. In this study, the masking ratio is fixed at *ρ* = 0.15, meaning that approximately 15% of the input features are randomly set to zero during each training pass. The corrupted input is computed as 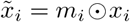, where ⊙ denotes element-wise multiplication.

The encoder receives the corrupted profile 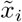, while the decoder is trained to reconstruct the original clean profile *x*_*i*_. Therefore, the reconstruction target is the complete omic vector, not the corrupted input. The reconstruction loss is computed as

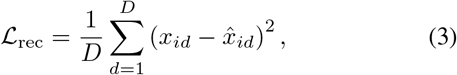

where 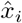 is the reconstructed output.

#### 3) Contrastive Representation Learning

Masked reconstruction encourages the latent representation to capture dependencies among omic features, such as genes. However, reconstruction alone does not explicitly align the latent space with survival outcomes. To introduce survival-aware geometry, CMVAE adds a supervised contrastive objective over confidently labeled samples.

First, we introduce a projection head *g*(·) that maps the deterministic latent mean into a normalized contrastive space:

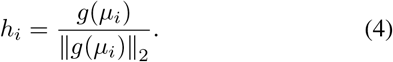

The contrastive objective is computed on *h*_*i*_, while *µ*_*i*_ serves as the final embedding used for classifier training and evaluation.

We adopt a supervised contrastive objective inspired by the similarity-based neighborhood preservation related to the soft-nearest-neighbor objectives [23], [24]. Let *B*_*L*_ denote the labeled subset of a mini-batch, and let *y*_*i*_ be the survival label for sample *i*. The temperature-scaled similarity between two projected embeddings is defined as: 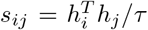, where *τ* is the temperature parameter (set to 0.2). For each labeled anchor sample *i*, samples from the same survival class form positive set, while samples from the opposite survival class act as negatives. In particular, the positive set is defined as:

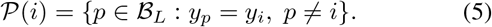

The supervised contrastive loss is computed as

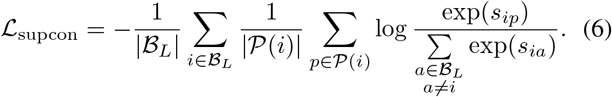

Only confidently labeled STS and LTS samples contribute to this term. Censored samples are excluded from Eq. 6, since they do not provide reliable survival-label relationships, but they remain active for the reconstruction loss.

#### 4) Objective Function

CMVAE is optimized using a joint objective consisting of masked reconstruction, variational regularization, and supervised contrastive learning as:

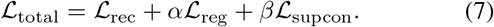

*β* controls the contribution of the supervised contrastive term which is kept small to ensure the contrastive objective refines the latent geometry without overpowering the reconstruction objective.

Two regularization variants are evaluated independently: Kullback-Leibler (KL) divergence [22] and Maximum Mean Discrepancy (MMD) [25]. The KL term is defined as:

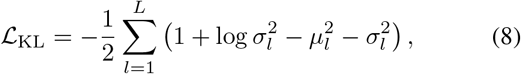

where *L* is the latent dimensionality. This term constrains the approximate posterior to remain close to the standard normal prior. The MMD regularization term matches the aggregate latent distribution to the standard normal prior:

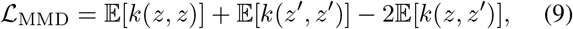

where *z* ∼ *q*_*ϕ*_(*z*|*x*), *z*^′^ ∼ *N*(0, *I*), and *k* denotes the RBF kernel. In our implementation, the maximum KL weight was set to 0.01, while the maximum MMD weight was set to 1.0. More about MMD and the selection of regularization coefficients are discussed in SI S1.

Overall, CMVAE learns from two complementary signals. The masked denoising objective forces the model to recover complete molecular profiles from corrupted inputs, while the supervised contrastive term explicitly imposes STS/LTS structure on the latent manifold. This allows the final representation to preserve both biological structure and survival-aware patient grouping.

### C. Baselines

We evaluate the performance of the proposed method against six distinct feature extraction and dimensionality reduction methods: Raw features (top 500 high-variance features), Principal Components (PCA) (top 50), Gini importance features (top 100), standard AEs, VAEs, and Masked Denoising VAEs (MVAEs) (details in SI S2). The deep generative models, i.e VAEs, MVAEs and CMVAEs are optimized with L2 reconstruction loss in combination with KL divergence and MMD regularization techniques independently. Additional VAEs are optimized using Logcosh reconstruction loss with the same two regularizers. The extracted features from each of these 12 techniques are subsequently fed into three machine learning classifiers: Logistic Regression (LR), Random Forest (RF), and Support Vector Machines with a polynomial kernel (PSVM). To rigorously handle class imbalance within the training pipeline, each classifier is evaluated under three distinct training configurations: a vanilla baseline with no over-sampling (NO), Random Oversampling (ROS), and Synthetic Minority Over-sampling Technique (SMOTE).

### D. Performance Evaluation Metrics

To rigorously assess and compare the predictive performance of the models, we utilize two primary evaluation metrics: the Area Under the Receiver Operating Characteristic curve (AUROC) [26] and the Concordance Index (C-index) [19]. Standard classification metrics, such as the *F*_1_-score and its maximum threshold variant (*F*_max_), are omitted due to their inherent limitations of heavy reliance on class prevalence. In highly imbalanced datasets, like ours, slight changes in the majority-to-minority ratio can distort these scores significantly which may lead to biased or misleading performance conclusions.

In contrast, AUROC and C-index offer robust, threshold-independent evaluations that are particularly well-suited for our experimental design for the survival prognosis prediction and stratification [12], [19]. The AUROC measures the model’s ability to discriminate between binary classes across all possible decision thresholds by plotting the true positive rate against the false positive rate. Complementing this, the C-index evaluates the rank correlation of predicted probabilities against true binary outcomes, quantifying the probability that the model correctly assigns a higher risk or likelihood score to a randomly selected positive instance compared to a randomly selected negative instance.

## III. Datasets

In this study, we focus only on the omics information and clinical outcomes relevant to the female Breast Invasive Carcinoma (BRCA). Since, BRCA is a highly heterogeneous malignancy with complex molecular and phenotypical classifications that drive diverse clinical outcomes [27], developing an accurate prognostic model would significantly benefit risk stratification after diagnosis.

We download four preprocessed single-omic datasets from the UCSC Xena Browser TCGA-BRCA cohort [28]^2^ representing different biological layers: transcriptomics (mRNA expression), epigenomics (DNA methylation), post-transcriptional regulation (miRNA expression), and genomics (copy number variation (CNV)). Clinical survival information is used to define short- and long-term survival (STS/LTS) labels.

For mRNA, we use the gene-level expression values in log_2_(*normalized count* + 1) format generated using the Illumina HiSeq platform. Since mRNA expression captures active transcriptional dysregulation, it provides a high-density molecular layer for learning survival-associated BRCA signatures. For epigenomic profiling, we use the DNA methylation values generated using the Illumina Infinium HumanMethylation450 platform. This represents as beta values, where each value ranges from 0 to 1 and reflects the relative methylation level at a CpG probe. DNA methylation is included because CpG-level methylation profiles capture regulatory and epigenetic changes that may contribute to BRCA survival differences. Next, we utilize the mature strand miRNA expression values, transformed as *log*_2_(*RPM* + 1), generated using the Illumina HiSeq miRNA-seq platform. Finally, We use the gene-level CNV dataset estimated using the GISTIC2 [29] method. CNV is included to evaluate whether large-scale genomic alteration patterns provide discriminative signal for BRCA STS vs. LTS prognosis. All four omics datasets are processed independently as single-omic cohorts rather than requiring complete patient overlap across modalities. For each modality, sample barcodes are matched with available TCGA-BRCA clinical survival metadata. The resulting modality-specific cohort comprises 1, 219 mRNA, 884 DNA methylation, 831 miRNA, and 1, 078 CNV samples.

### A. Short-& Long-term Sample Labeling: STS vs. LTS

This work correspond to only the uncensored patients in survival analysis where the patients exact time-to-event (such as the date of death, disease progression, or recovery) is fully known and recorded. The censored patients are not used in the contrastive learning, classifier training and performance analysis. We label patients STS and LTS based onfive-years survivability. Because the overall survival time (OS.time) in the dataset is recorded in days, we define the five-year threshold as 1826.25 days. Our main goal is to identify patients who do not survive beyond the five-year mark. Therefore, we designate STS as the positive class (label 1) and LTS who lived past five years as label 0. Patients who are still alive but have a follow-up time of less than five years are excluded from the analysis, since their ultimate survival category cannot yet be determined. Fig. 2 shows the highly imbalanced label distribution divided into omics types where the number of STS falls significantly short of LTS. Notably, significant number of samples are censored, thus cannot be used in classifier evaluation.

**Fig. 2.**
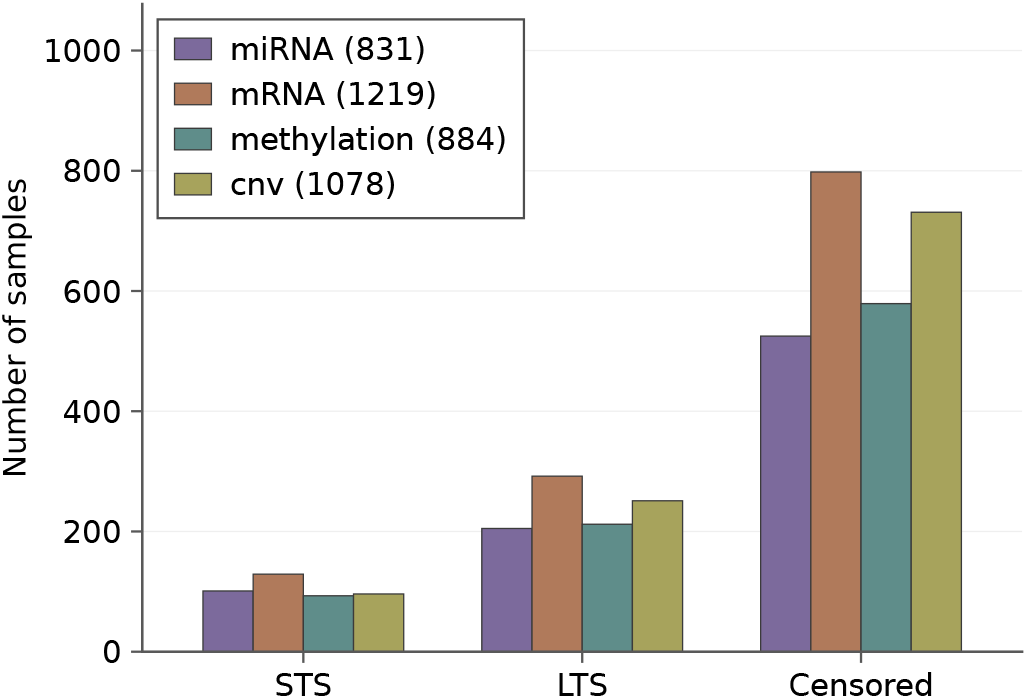
Label distribution of the TCGA Breast Invasive Carcinoma (BRCA) cohort across four omics layers.

## IV. Results and Discussion

To analyze the effectiveness of our proposed single-omic-based CMVAE, we evaluate and compare on both patient prognosis and survival stratification tasks. Beyond, we compared with multi-omics integration methods.

### Biological signals are enhanced most by contrastive supervision in transcriptomics for prognosis (Fig. 3A)

The CMVAE-based classifiers significantly outperform all baselines, while the baselines are better than the random cointoss classifier for mRNA (Fig. 3A). Specifically, CMVAE variants across both objectives and sampling techniques achieve AUROCs of 0.71 − 0.74 (Fig S1A). This highlights the ability of CMVAE to capture distinct transcriptional dysregulations in BRCA molecular subtypes, such as Basal-like and Luminal groups, and clinical outcomes in latent representation. In contrast to that, vanilla AEs and VAEs tail behind the linear variance-based dimensionality reduction methods, such as PCA and raw features which achieve a minimum of 0.63 to maximum of 0.72 AUROC across all the variants of machine learning models (SI Fig S1A). This indicates that global variance in gene expression remains a strong predictor of clinical survival and that contrastive learning effectively structures the latent manifold to reflect these outcomes.

**Fig. 3.**
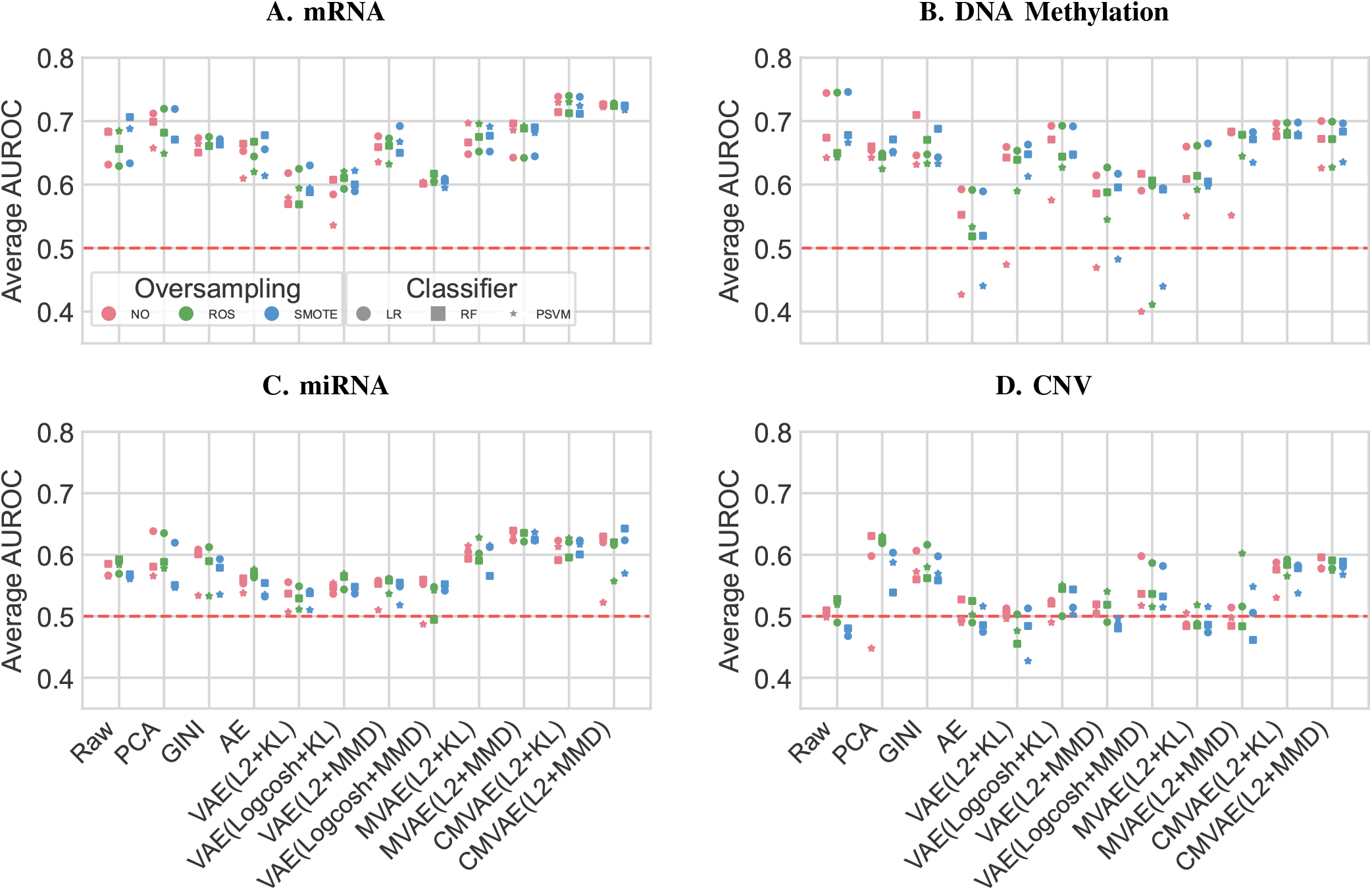
Performance evaluation comparison in classifying STS vs. LTS BRCA patients across 12 dimensionality reduction methods in four omics layers: (A) mRNA, (B) DNA Methylation, (C) miRNA, and (D) CNV. CMVAE learns clinical outcome aware representations across all omics types.

### CMVAE learns consistent and stable features across all four omics types (Fig. 3 and 4)

CMVAE preserves prognostic signatures that are lost by unsupervised vanilla reconstruction objectives by jointly optimizing omics with similar survival outcomes while separating divergent patients. In addition to mRNA expression, contrastive frameworks achieve superior and consistent performance when comparing all three classifiers for DNA methylation, while the raw top 500 most variant features yields the highest discriminative power. These raw representations outperform latent variable modeling, achieving a AUROC of 0.7459 ± 0.0911 compared to the CMVAE’s peak AUROC of 0.7001 ± 0.1027 (Fig. 3B and SI Fig S1B). Since these uncompressed feature sets preserve hypermethylation markers in promoter regions, they provide strong biological signals to simple linear classifiers. This aligns with the remark that linear classifiers frequently outperform deep neural networks on highly sparse methylation profiles [30].

**Fig. 4.**
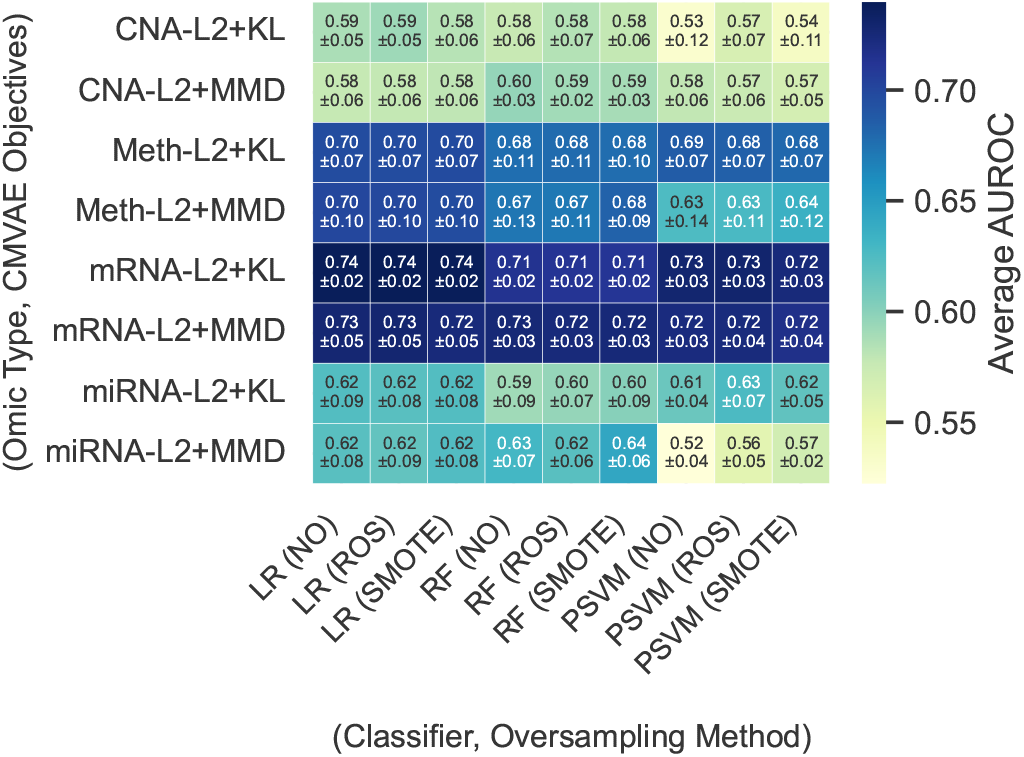
Detailed performance of CMVAE across the combination of three classifiers with three oversampling techniques for BRCA patient prognosis.

Even within inherently weaker modalities, the contrastive loss demonstrated its robustness by successfully stabilizing and extracting non-random prognostic patterns. In the miRNA cohort, the contrastive approach achieves clear performance gains, enabling an L2 + MMD objective paired with a Random Forest (RF) classifier to achieve a peak AUROC of 0.6426 ± 0.0614. For the gene-level CNV data that carries the most sparse biological signals and underperforms in standard architectures, the contrastive framework enforces noticeable improvements and stabilizes the latent space, yielding a peak AUROC of 0.5958± 0.0364 (L2 + MMD, RF) (Fig. 4 and SI Fig S1C,D).

Across all four omic types, CMVAE is the most effective dimensionality reduction strategy. While the baseline signal abundance varied by omic type, the supervised contrastive strategy universally enhanced the network’s capability to isolate and protect critical clinical signatures during data reconstruction. They optimally extract relevant biological signals for patient prognosis and STS prediction across all three evaluated machine learning classifiers.

### Survivability analysis: Kaplan-Meier (KM) curve (Fig. 5)

To evaluate whether classifier-derived CMVAE risk scores correspond to actual patient survival patterns, we perform the KM survival analysis on the BRCA mRNA cohort. The best-performing CMVAE configuration is first selected using the mean AUROC across the five cross-validation folds. For each fold, the saved fold-specific preprocessor, trained CMVAE encoder, and classifier were utilized without retraining. The held-out validation samples from that fold are transformed using the corresponding training-fold preprocessing pipeline, encoded into the CMVAE latent space, and passed to the trained classifier to obtain an out-of-fold predicted STS probability. Thus, each patient receives a risk score from a model that was not trained on that patient.

**Fig. 5.**
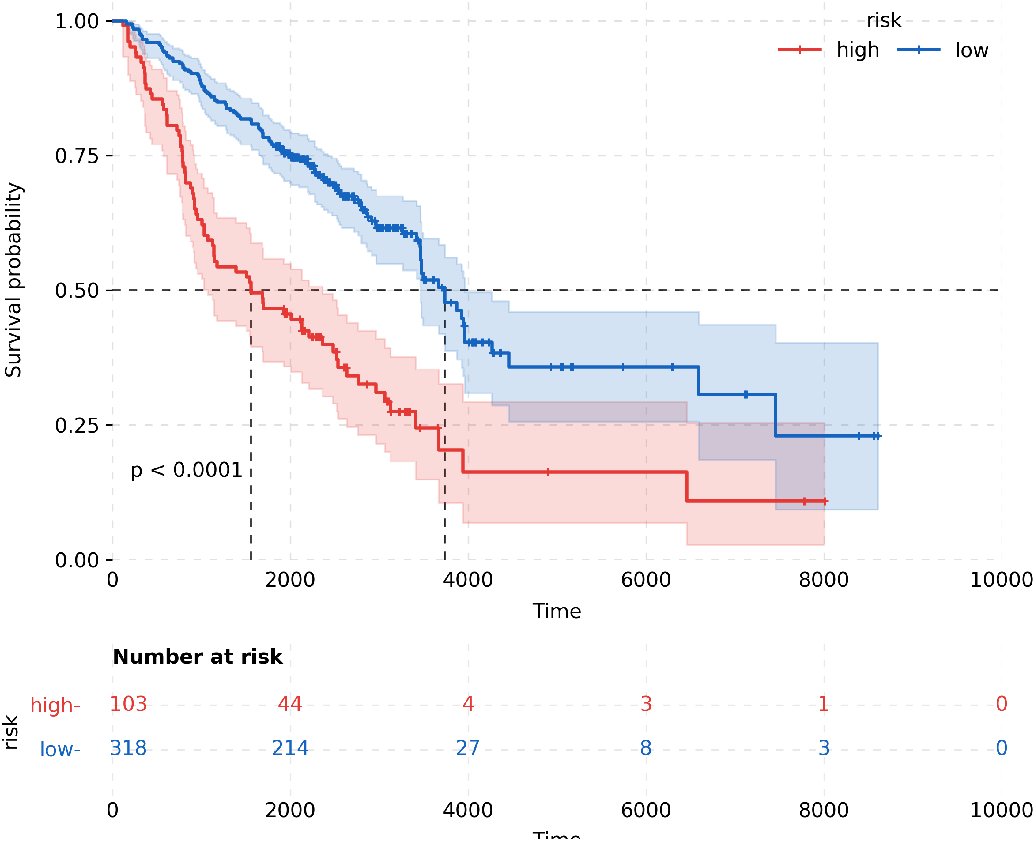
Kaplan-Meier (KM) survivability analysis between two BRCA risk groups.

The predicted STS probability is used as the model-derived survival risk score. Patients with predicted STS probability greater than or equal to 0.5 are assigned to the high-risk group, while patients below 0.5 are assigned to the low-risk group. KM curves are then fitted using the true overall survival time (*OS*.*time*) and event indicator (*OS*) for these two predicted risk groups. Censored patients are retained in the KM estimation through the event indicator and marked on the curve, while the number-at-risk table is computed at the displayed follow-up time points. A log-rank test is also performed between the high- and low-risk groups to quantify whether their survival distributions are significantly different.

KM analysis based on the BRCA mRNA cohort demonstrates that the CMVAE-derived risk score separates patients into prognostically distinct survival strata (Fig. 5). We observe a clear separation between the high- and low-risk groups, supported by a log-rank significance of *p <* 0.0001. The low-risk cohort maintains consistently higher survival probability across the follow-up period, whereas the high-risk cohort exhibits a more rapid decline. This indicates earlier event occurrence among patients assigned higher predicted STS risk. Because the risk groups were derived from out-of-fold model predictions, this separation suggests that CMVAE learns a latent representation with clinically meaningful survival structure rather than simply memorizing the training data. This analysis complements AUROC-based classification results by showing that the learned mRNA latent space is not only discriminative for five-year STS/LTS classification, but also capable of producing risk strata with significantly different overall survival trajectories.

### Concordance index (C-index) analysis and comparison with multi-omics integration methods (Fig. 6)

To further evaluate whether the learned transcriptomic (mRNA) representations preserve survival ordering beyond a binary setup, we compute Harrell’s C-index [19] using the out-of-fold classifier-derived STS risk scores. The C-index evaluates whether patients with higher predicted STS risk scores also show earlier observed survival events. For this study, since the classifier outputs are a probability for the STS class, the larger values would indicate higher chances of short-term survival risk, which means shorter expected survival. Hence, a negative STS risk score is used as the predicted survival score for concordance calculation. The CMVAE achieves a mean fold-level C-index of 0.688 ± 0.031, with a pooled out-of-fold C-index of 0.689 and a bootstrap 95% confidence interval of [0.649,0.730] (Fig. 6A). This indicates that the learned risk score ranks patient survival outcomes substantially above random concordance, supporting the presence of survival-relevant structure only on the confidently labels STS/LTS cohort using the held-out fold.

**Fig. 6.**
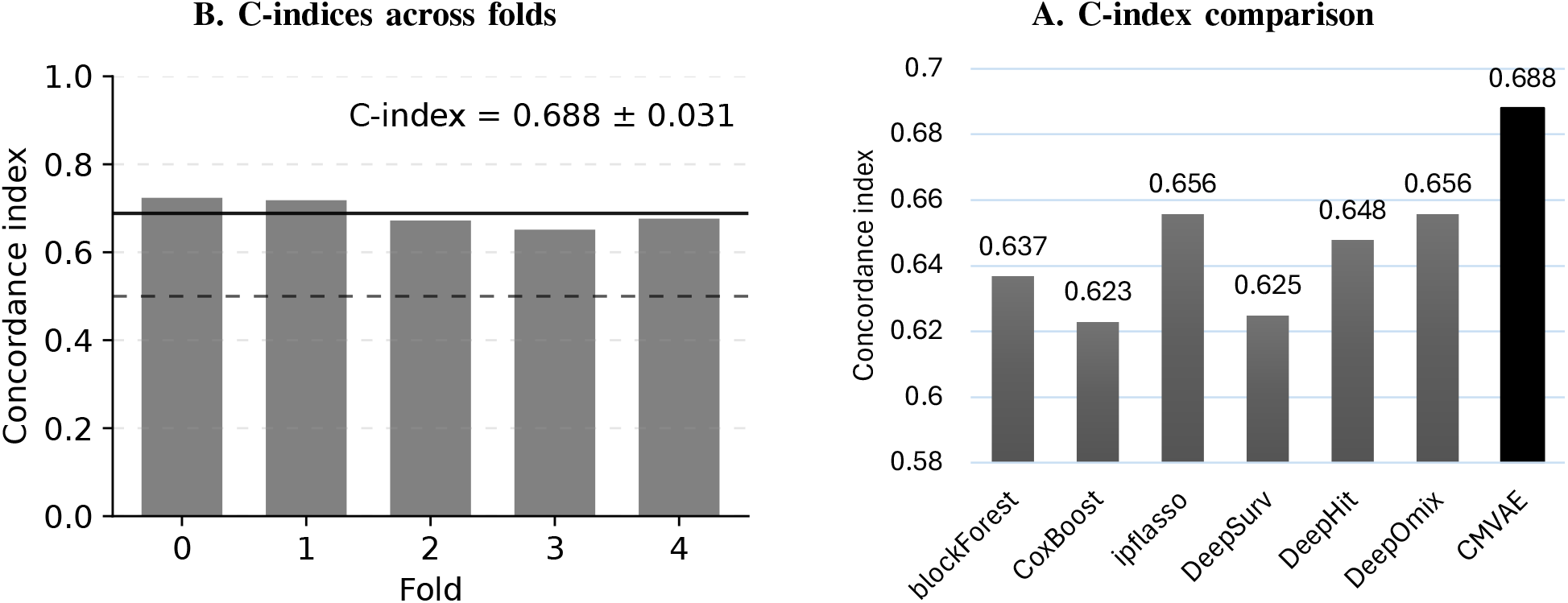
Concordance-index (C-index) analysis. (A) Consistent C-index values across stratified folds. Dotted and continuous line denote the random baseline and mean C-index, respectively. (B) C-index comparison with six multi-omics methods.

C-index comparison with existing multi-omics and deep survival approaches further demonstrates the prognostic strength of the proposed CMVAE framework (Fig. 6B). CMVAE achieves the highest concordance index of 0.688, outperforming blockforest (0.637), CoxBoost (0.623), ipflasso (0.656), DeepSurv (0.625), DeepHit (0.648), and DeepOmix (0.656) (sourced from DeepOmix [12]). This indicates that the learned contrastive transcriptomic representation provides stronger survival ranking than both classical and multiomics survival models and deep survival architectures. The improvement over DeepSurv [31] and CoxBoost [32]–[34] further suggests that the CMVAE latent space captures prognostic transcriptomic structure that is not fully recovered by standard survival modeling frameworks.

### Discussion: Reconstruction objectives may capture input manifold, but without implicitly encoding prognostic outcomes

Pure reconstruction objectives in standard AEs and VAEs inherently prioritize capturing global input manifolds over isolating specific prognostic signatures. Consequently, experimental results across all omic modalities demonstrate that unsupervised reconstruction, regardless of the regularization technique applied, fails to effectively encode clinical survival boundaries in the latent space (Fig. 3, SI Fig. S1). Although masked denoising variants of VAEs (MVAE) offer marginal performance and stability gains by mitigating technical noise, they consistently underperform or merely match simpler linear baselines, such as PCA. In signal-dense modalities such as mRNA and DNA methylation, MVAEs plateaued at peak AUROCs of approximately 0.69 and 0.68, respectively, falling short of standard PCA and raw feature baselines (AUROCs of 0.72–0.75). For inherently weaker modalities, miRNA and CNV, the unsupervised framework provided minimal utility. Collectively, these findings underscore that optimizing for data reconstruction is fundamentally misaligned with downstream clinical classification tasks. Without a supervised or alignment mechanism, unsupervised VAEs leave prognostic predictions within a clinically insignificant range.

## V. Conclusion and Future Work

We presented CMVAE, a dual-optimization-based dimensionality reduction method that addresses both the low patient overlap across multi-omics data and posterior collapse in pure reconstruction objectives. By optimizing a joint loss function that combines masked reconstruction and a supervised contrastive objective, CMVAE learns a clinical outcome-aware latent representation space by preserving the molecular manifold.

Rigorous evaluation within a leak-proof five-fold stratified cross-validation pipeline and KM survival analysis in the TCGA-BRCA cohort confirms that CMVAE is effective on both predicting clinical outcomes and high-risk patient stratification. In single-omic evaluations, CMVAE out-performed standard unsupervised baselines with a peak AUROC of 0.7397 ± 0.0239 on mRNA data. Downstream stratification analysis demonstrates a highly significant log-rank differentiation between low- and high-risk patient subgroups. Moving beyond single-omic inputs, CMVAE demonstrates robust superiority in integrative multi-omics tasks, outperforming six state-of-the-art multi-omics survival baselines.

Future efforts will aim to expand the generalizability of this model by performing pan-cancer validations across diverse TCGA cohorts to evaluate its robustness. Additionally, we plan to leverage post-hoc explainability methods to isolate specific driving cross-omic biomarkers. Translating this architecture into a deployable clinical decision-support tool will ultimately require rigorous external validation on independent, prospective clinical trial data to ensure its real-world utility. Finally, a natural direction for expanding this paradigm will focus on developing multi-view, task-agnostic cross-omic representation learning, enabling the model to serve as a foundational multiomics embedding framework.

## Supplementary Information

### S1. Maximum Mean Discrepancy (mmd) and EFfective Coefficient Selection

The MMD regularization term matches the aggregate latent distribution to the standard normal prior:

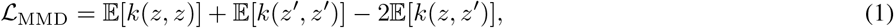

where *z* ∼ *q*_*ϕ*_(*z*|*x*), *z*^′^ ∼ *N* (0, *I*), and *k* denotes the RBF kernel. The empirical estimate of the first expectation is

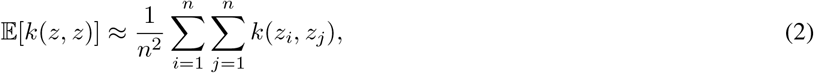

where 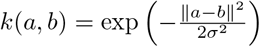, *σ* = 1.0.

Regularization coefficient: To reduce posterior collapse, the regularization coefficient is linearly warmed up during early training. For epoch *t*, the ramp-up factor is 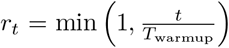 . The effective KL weight is computed as *β*(*t*) = *r*_*t*_*β*_max_, and the effective MMD weight is computed as *λ*(*t*) = *r*_*t*_*λ*_max_.

### S2. Baseline Implementation Details

#### A. Uncompressed Baseline: Top 500 Highest-Variance Raw Features

To prevent data leakage, we compute preprocessing parameters strictly within each training fold and apply them deterministically to the corresponding validation fold. Top 500 highest-variance omic features are kept exclusively from the training partition to eliminate uninformative noise. Missing values are then imputed using feature-wise training medians. Remaining features are *z*-score standardized using scaling parameters fit solely on the training fold. We keep the same exact 500 features from validation fold, apply the training fold medians and the *z*-score scaling on the validation set. This ensures no-leakage from training to validation sets and gives an unbiased evaluations across classifiers.

#### B. PCA-Reduced Space: 50 Principal Components via Training-fold-Isolated Projection

To isolate feature extraction and eliminate data leakage, a fold-specific PCA pipeline is implemented. Features with *>* 20% missing or 0-values are filtered out. Missing entries are filled using training-fold feature medians, and data was z-score standardized exclusively on training data to balance feature magnitude. The learned parameters are then applied to transform validation fold. Next, PCA is fit on the standardized training partition, retaining the top 50 principal components. The learned projection matrix is applied independently to the validation fold. The 50-dimensional representations served as inputs for the downstream classifiers (LR, RF, Poly-SVM). For the SMOTE condition, synthetic sample generation is performed directly within this reduced space to improve the geometric validity of the interpolations.

#### C. Supervised Feature Baseline: Top 100 Features via Training-Isolated Gini Importance

To isolate supervised feature selection and prevent class-imbalance bias, a fold-independent Gini importance pipeline is implemented. Features with *>* 20% missing or zero entries are filtered out. Remaining missing values are filled via training feature medians, and the data is normalized using a MinMaxScaler fit solely on the training fold. A Random Forest selector (100 estimators) is trained on the scaled training partition. Balanced class weights are applied to counteract the severe 5-year survival class imbalance in the patient cancer cohort, ensuring unbiased importance scores. Features are ranked by mean decrease in Gini impurity, and the top 100 omic feautres are extracted. The selected feature indices are sorted to preserve original structural positioning. Minority oversampling (ROS, SMOTE) is confined strictly to this 100-dimensional training subspace, ensuring the feature selector operated without any exposure to the entire dataset or validation folds.

#### D. Auto Encoder (AE)

In this workflow, we perform dimensionality reduction using a deterministic autoencoder (AE), trained independently within each cross-validation fold to prevent data leakage. Input omic data underwent zero and NaN preprocessing, followed by train-fold median imputation and *z*-score standardization fitted exclusively on the training sets.

The symmetric AE architecture compresses features through three fully connected encoder layers (512 → 256 → 128) into a 32-dimensional latent space via a linear projection, which the decoder mirrors in reverse. All hidden layers utilize rectified linear unit (ReLU) activations and a 0.3 dropout rate. The final decoder layer employs a linear activation to reconstruct the standardized inputs. To mitigate overfitting given the high-dimensional, low-sample-size data, the network is optimized using Mean Squared Error (MSE) loss, the Adam optimizer (learning rate = 10^−3^), and *L*_2_ weight regularization (weight decay = 10^−5^). Each training fold was internally partitioned into an 85/15 split. The internal validation MSE was monitored for early stopping (epochs = 50, patience = 15, threshold = 10^−6^), and the best-performing weights were restored post-training. To ensure independent initialization and reproducibility, seeds were determined via ae seed = SEED + fold. The CV holdout fold was isolated entirely from training and early stopping, eliminating indirect data leakage.

Following optimization, the trained encoder extracted 32-dimensional latent representations for both training and validation folds. To optimize computational efficiency, these embeddings were cached per fold and reused across all downstream classifier configurations. To accommodate the varying scale-sensitivity of the models, two variants of the latent space were prepared: *z*-score normalized embeddings—scaled via a `StandardScaler` fitted exclusively on the training fold’s representations—were provided to Logistic Regression (LR) and polynomial Support Vector Machine (poly-SVM) classifiers. Conversely, Random Forest (RF) classifiers received raw, unscaled encodings due to the scale-invariance of tree-based splitting criteria.

Class imbalance was managed using three strategies: No Oversampling (NO), Random Over-Sampling (ROS), and SMOTE. In the NO condition, the `class_weight=‘balanced’` parameter was applied; this parameter was mutually exclusive with oversampling to prevent redundant correction of the loss landscape. For SMOTE, the number of nearest neighbors was dynamically bounded at min(5, *n*_minority_ − 1) to accommodate small minority classes. All classifiers—LR (max iter = 1000), RF (n estimators = 100), and poly-SVM (degree = 3, probability = True)—were deterministically initialized per fold using SEED + fold to guarantee reproducibility.

#### E. Variational Autoencoder (VAE) Pipeline and Architecture

The Variational Autoencoder (VAE) pipeline utilized identical zero and NaN miRNA preprocessing, train-fold median imputation, and *z*-score standardization as the deterministic AE pipeline. Preprocessing was executed once per cross-validation (CV) fold and shared across all four evaluated loss configurations. This ensures that downstream classifier performance variations are strictly attributable to the VAE training objectives rather than input discrepancies.

The network employs an asymmetric encoder-decoder architecture. The encoder features three shared fully connected hidden layers (input → 512 → 256 → 128), each utilizing ReLU activations and a 0.3 dropout rate. The final hidden layer maps to two parallel linear projection heads (*µ* and log *σ*^2^) that parameterize the approximate variational posterior distribution by mapping ℝ^128^ → ℝ^32^.

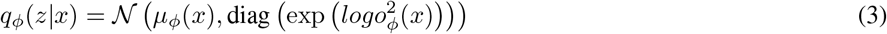

Where *µ*_*ϕ*_(*x*) and 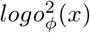 are the outputs of *fc_mu* and *fc_log_var*.

Conversely, the decoder maps the 32-dimensional latent space back to the original input dimensionality via three symmetric hidden layers (128 → 256 → 512 → input) and a final linear output layer, matching the *z*-score standardized input format under a Mean Squared Error (MSE) reconstruction loss. Gradient flow through the stochastic latent node was enabled during training via the reparameterization trick.

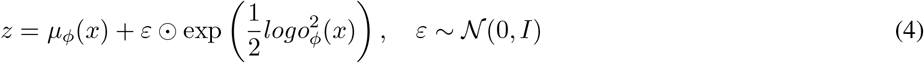

At inference time, the stochastic sampling step is omitted. Downstream classifiers receive only the deterministic posterior mean, *µ*_*ϕ*_(**x**). This eliminates sampling noise, ensuring the generation of stable, reproducible feature vectors from each sample’s omic profile.

#### F. Masked Variational Autoencoder (MVAE) Pipeline and Architecture

The primary representation learning framework in this study was designed around a masked denoising variational autoencoder (MVAE). The motivation for this architecture was that high-dimensional omic profiles contain correlated molecular signals, but direct reconstruction of the full input can allow a neural network to learn a near-identity mapping without necessarily extracting robust biological structure. To address this, the MVAE was trained to reconstruct the original, complete omic profile from a partially corrupted version of the same sample. This forces the encoder to infer missing molecular information from the remaining observed features and encourages the latent space to capture dependencies across the omic profile.

### S3. Performance Comparison

Fig. S1 reports the average performance gains (AUROC) and standard deviations for all methods evaluated via leak-proof, five-fold stratified cross-validation in BRCA patient prognosis. These approaches comprise three machine learning classifiers; each classifier is paired with three oversampling techniques using features extracted from seven distinct dimensionality reduction methods, including the proposed CMVAE.

**Fig. S1.**
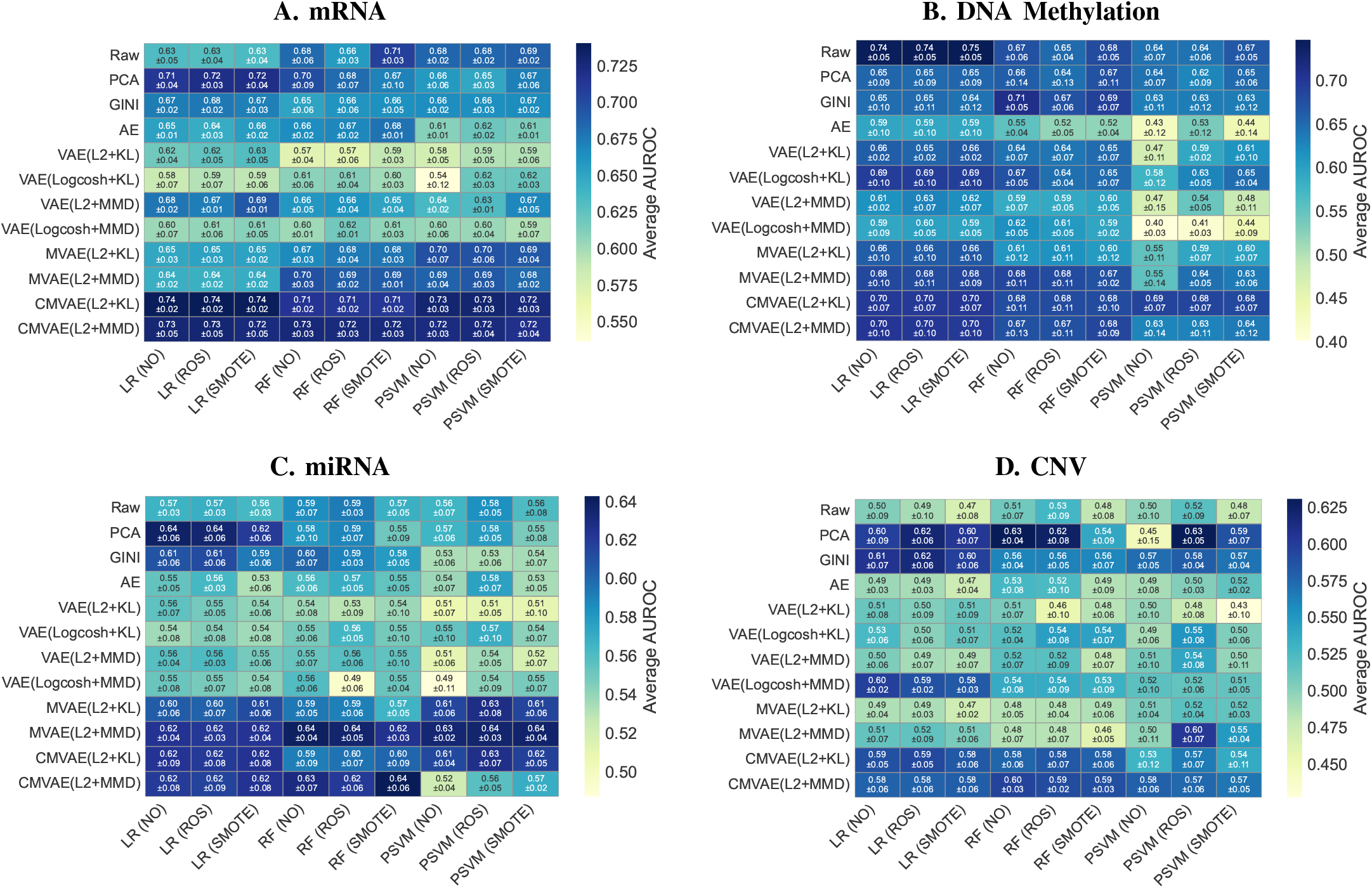
Performance comparison for classifying short-vs. long-term survival in BRCA patients across four omic modalities: (A) mRNA, (B) DNA methylation, (C) miRNA, and (D) CNV.

## Footnotes

1 This work has been accepted at the IEEE BIBM 2026 workshop RLPDBio (Representation Learning and Pattern Discovery for Bioinformatics and Biomedicine).

2 https://xenabrowser.net/datapages/

